# Deep Batch Active Learning for Drug Discovery

**DOI:** 10.1101/2023.07.26.550653

**Authors:** Michael Bailey, Saeed Moayedpour, Ruijiang Li, Alejandro Corrochano-Navarro, Alexander Kötter, Lorenzo Kogler-Anele, Saleh Riahi, Christoph Grebner, Gerhard Hessler, Hans Matter, Marc Bianciotto, Pablo Mas, Ziv Bar-Joseph, Sven Jager

## Abstract

A key challenge in drug discovery is to optimize, in silico, various absorption and affinity properties of small molecules. One strategy that was proposed for such optimization process is active learning. In active learning molecules are selected for testing based on their likelihood of improving model performance. To enable the use of active learning with advanced neural network models we developed two novel active learning batch selection methods. These methods were tested on several public datasets for different optimization goals and with different sizes. We have also curated new affinity datasets that provide chronological information on state-of-the-art experimental strategy. As we show, for all datasets the new active learning methods greatly improved on existing and current batch selection methods leading to significant potential saving in the number of experiments needed to reach the same model performance. Our methods are general and can be used with any package including the popular `DeepChem` library.

## Introduction

The process that leads from a small molecule that shows some activity against the target of interest to a candidate for clinical development involves complex multi-parameter optimization. In this process, in addition to the activity against the target itself, the ADMET profile of the molecule, i.e. its Absorption, Distribution, Metabolism, Excretion, and Toxicity properties are optimized. Accurate *in silico* models for the desired properties are required to speed up and improve decision making and reduce the number of necessary experiments***Hessler and Baringhaus (2018); Grebner et al. 2022); Wu et al. (2020)***. Amongst other machine learning (ML) techniques, deep learning models, and more specifically (graph) neural neural networks, have been used successfully in this field***Xiong et al. (2021)***.

Scale-effective ADMET and affinity prediction methods require an abundance of labeled training data due to their complexity and the need to cover the enormous molecular design space. This can be a bottleneck in cost, time, and experimental resources***Chuang et al. (2020)***. Various approaches have been proposed for improving data acquisition and the models, including transfer learning ***Weiss et al. (2016)***, data augmentation ***Cai et al. (2020)*** and *active learning* ***Cohn et al. (1996a)***.

Current optimization approaches often work in cycles. In each cycle a set of molecules are tested, the model is revised and, based on the revised model, a new set is selected for testing, and so on until the model reaches the desired performance.

Active learning ***Cohn et al. (1996b)*** is an approach for selecting molecules for each of the cycles. Unlike traditional approaches that test the most promising candidates in each round ***Cohn et al. (1994)*** in active learning samples are selected to optimize the *model* rather than the cycle result. In such approach optimization samples are prioritized by their ability to improve model performance when labeled. Active learning has been studied in sequential mode, where samples are labelled one-at-a-time,and batch mode, where samples are selected for labelling in batches ***Settles (2012)***. Batch mode is both more realistic for small molecule optimization and more challenging computationally. The main problem is that samples are not independent (sharing chemical properties that will influence model parameters) and so selecting a set based on marginal improvement does not reflect well the improvement provided by the entire batch ***Ash et al. (2021)***.

Active learning methods have been utilized to predict and optimize the physiochemical and biological properties of molecular systems. For instance, batch active learning (BMDAL) was used to enhance the model accuracy in predicting the conformations, energetics, and interatomic forces of small organic compounds where the diversity of the training data is a limiting factor.***Zaverkin et al. (2022)***. Active learning was also combined with pairwise difference regression (PADRE)***Tynes et al. (2021)*** to predict molecular properties including redox free energy. Reker et al developed an active learning based tool to screen and select top candidates for ligand-target binding prediction ***Reker et al. (2017)***. Thompson *et al*.***Thompson et al. (2022)*** employed Active learning framework to build a package for the binding free energy of to TYK2 Kinase. In another study model uncertainty for unlabeled pharmacokinetics was used to set an active learning pipeline for plasma exposure prediction ***Ding et al. (2021)***. Pertusi et al ***Pertusi et al. (2017)***, used an active learning data selection method for characterizing enzyme promiscuity. Naik et al ***Naik et al. (2016)*** used active learning to build models to predict molecule effects on subcellular protein localization. However, while some of these methods worked well, they were not applied to the more advanced deep learning methods that have become the tool of choice for small molecule modeling.

On the theoretical side, a number of Batch Active learning methods have recently been developed though these have not been used in the drug design space. For example, BAIT***Ash et al. (2021)*** uses a probabilistic approach for the learning procedure that optimally selects (using greedy approximation) a set of samples that maximizes the likelihood of the model parameters (last layer) as defined by Fisher information***Ash et al. (2021)***. Others have proposed using the local approximation to estimate the maximum of the posterior distribution over the batch. This is achieved through computation of the inverse Hessian of the negative log posterior ***Daxberger et al. (2021)***. Recently, GeneDisco***Mehrjou et al. (2021)*** was published as an open source library of benchmarking data pertaining to transcriptomics active learning work. However, these methods have not been extensively tested for small molecules optimization. Specifically, to date, popular *in silico* design suits, including ChemML ***Haghighatlari et al. (2019)*** and DeepChem ***dee (2016)***, do not support active leaning strategies.

To address these shortcoming, we developed a new innovative and generalizable strategy that can be used with any deep learning ADMET methods. Our methods are inspired by the Bayesian deep regression paradigm, where estimating the model uncertainty is tantamount to obtaining the posterior distribution of the model parameters***Kendall and Gal (2017)***. Model uncertainty is determined using innovative sampling strategies, with no extra model training required.We next select batches that maximize the joint entropy, i.e., the log-determinant of the epistemic covariance of the batch predictions. This enforces batch diversity by rejecting highly correlated batches.

To evaluate our methods and compare them to state of the art approaches we have assembled a large collection of datasets. As we show, our active learning algorithm consistently leads to the best performance when compared to prior methods suggested for this task.

We also generated new data for internal candidates and show that our methods can significantly save cost and time compared to current industry optimization approaches for these datasets.

## Results

We developed and tested several batch active learning selection approaches which quantify the uncertainty over multiple samples. Given these uncertainties our method aims to select the subset of samples with maximal joint entropy (i.e., information content) (Figure 1). Specifically, we use multiple methods to compute a covariance matrix, *C*, between predictions on unlabeled samples, 𝒱. Then, using an iterative and greedy approach, the method selects a submatrix *C*_*B*_ of size *B* × *B* from *C* with maximal determinant. Such approach takes into account both the “uncertainty” (which is manifested in the variance of each sample) and the “diversity” (which is reflected in the covariance). See Methods for details.

**Figure 1.**
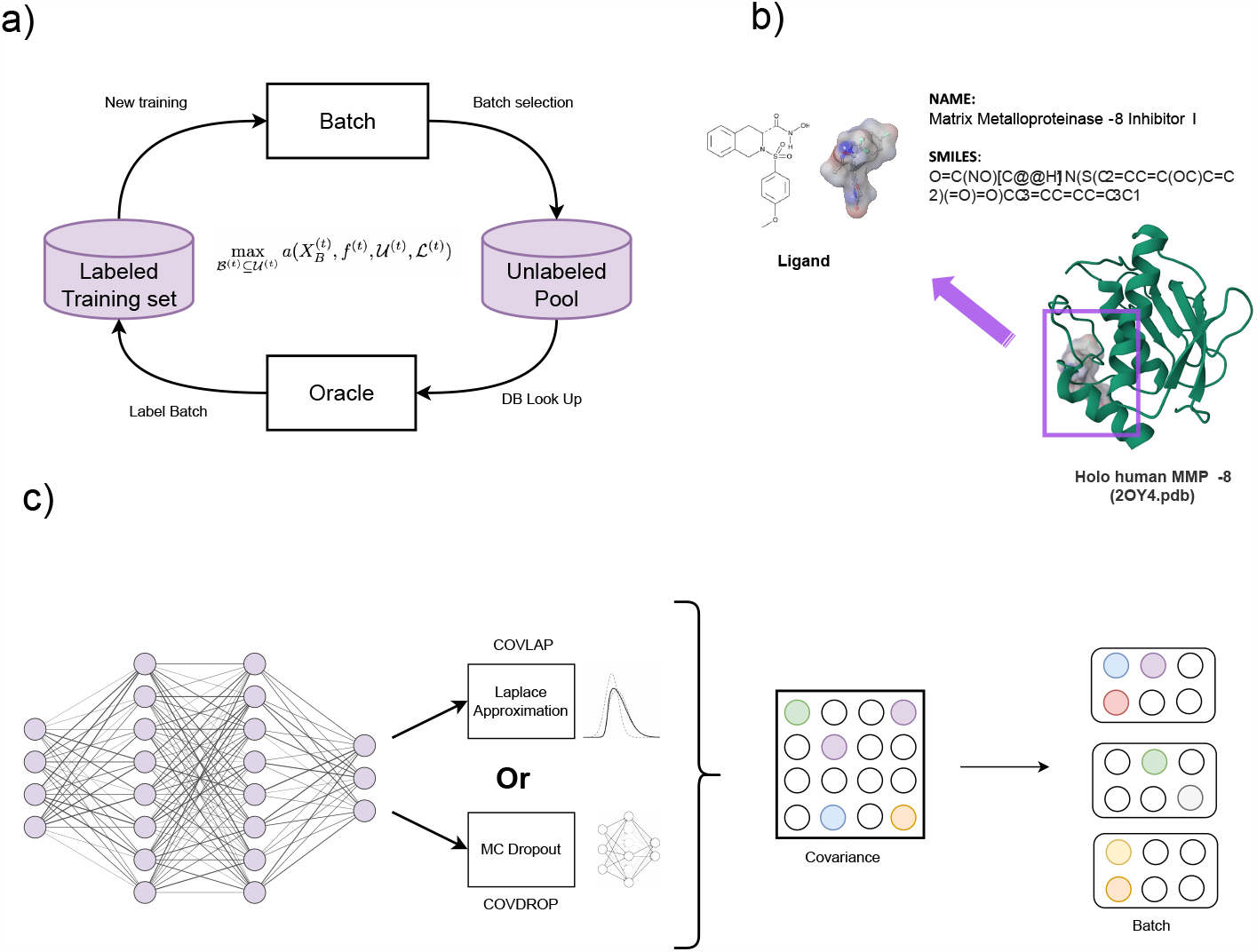
Method overview. a) The active learning loop. We consider the entire dataset as the pool, and the oracle as the entity that holds the corresponding labels. During each iteration of the active learning loop, we select a batch of points from the pool and obtain their corresponding labels from the oracle. We then update our model using the selected batch and evaluate its performance on a holdout set. This process is repeated until the desired level of performance is achieved. b) Prediction of binding affinity is the target function for the ChEMBL and Sanofi-Aventis datasets c) Active learning batch selection. At the last layer of our model, we use either Laplace approximation or Monte Carlo dropout to compute covariances (COVLAP and COVDROP), from which an ensemble of predictions is generated. With the derived covariance matrix, we optimize batches iteratively based on their information content.

### Evaluating active learning method on ADMET and affinity related data

We used several public drug design datasets to test and compare the performance of our methods. For comparison, in addition to the two methods we developed (MC dropout and Laplace Approximation, COVDROP and COVLAP, respectively) we also compared to methods that have been previously suggested (*k*-means***Nguyen and Smeulders (2004)*** and BAIT***Ash et al. (2021)*** and to a random ordering of the experiments (i.e. no active learning). Batch size was set to 30 for all methods. During each iteration of the loop, each model (e.g., *k*-Means, BAIT, Random, COVDROP, or COV-LAP) selects a batch consisting of a fixed number of samples from the unlabeled pool. This iterative process is repeated until all labels in the Oracle are exhausted. In the retrospective setting, the pool includes all samples from the relevant dataset, while the oracle retains the corresponding labels.

Our evaluation datasets included a cell permeability dataset with 906 drugs ***Wang et al. (2016)***, aqueous solubility dataset, which comprises 9,982 small molecules ***Sorkun et al. (2019); Huang et al. (2022)***, and the lipophilicity data for 1200 small molecules ***Wenlock and Tomkinson (2015); Wu et al. (2018)***. We also included 10 large affinity datasets, 6 from ChEMBL and 4 new internal datasets. Details for all datasets are provided in Methods and in Table 1.

**Table 1.** Number or required experiments for reaching model error with 10% higher than the minimum calculated RMSE for the whole training set across different methods. NC - Number of compounds in this dataset. Chron - analysis performed with batch selection using the actual order in which the data was profiled (only available for internal data). % gain estimates the improvement over the Random selection computed from (1 − N_COVDROP_/N_Random_) × 100. Where N_COVDROP_ as well as N_Random_ represents the average number of experiments out of 25 active learning cycles executed and a batch size of 32.

| Dataset | COVDROP | COVLAP | k-means | BAIT | Random | Chron. | NC | % gain |
| --- | --- | --- | --- | --- | --- | --- | --- | --- |
| Caco2 | 420 | 450 | 570 | 540 | 600 | - | 637 | 30.0 |
| Lipophilicity | 1440 | 1500 | 1560 | 1650 | 1650 | - | 2940 | 12.7 |
| HFE | 360 | 450 | 450 | 450 | 450 | - | 450 | 20.0 |
| Solubility | 2160 | 6810 | 5850 | 6988 | 5730 | - | 6988 | 62.3 |
| PPBR | 1050 | 450 | 450 | 420 | 510 | - | 1130 | - |
| GSK3β | 1260 | 2325 | 2325 | 2325 | 2325 | - | 2325 | 45.8 |
| MMP3 | 690 | 810 | 900 | 870 | 930 | - | 1280 | 25.8 |
| PPARδ | 570 | 660 | 660 | 630 | 690 | - | 879 | 17.4 |
| A1AR | 570 | 540 | 600 | 630 | 630 | - | 952 | 9.5 |
| NAV17 | 1770 | 4004 | 4004 | 4004 | 4004 | - | 4004 | 55.8 |
| D2 | 2010 | 4797 | 4797 | 4797 | 4797 | - | 4797 | 58.1 |
| Renin | 150 | 240 | 270 | 300 | 330 | 390 | 422 | 54.5 |
| FXa | 750 | 1020 | 870 | 1026 | 990 | 1026 | 1026 | 24.2 |
| MMP-8 | 150 | 270 | 390 | 420 | 456 | 360 | 456 | 67.1 |
| PPARδ | 420 | 300 | 570 | 390 | 570 | 600 | 614 | 26.3 |

### Comparison of active learning methods for solubility datasets

Results for a selected subset of the benchmarked datasets are presented in Figure 2. In these figures we present the accuracy of models when using the different active learning methods as a function of the iteration. As can be seen, in most cases the COVDROP method very quickly leads to better performance when compared to other methods.

**Figure 2.**
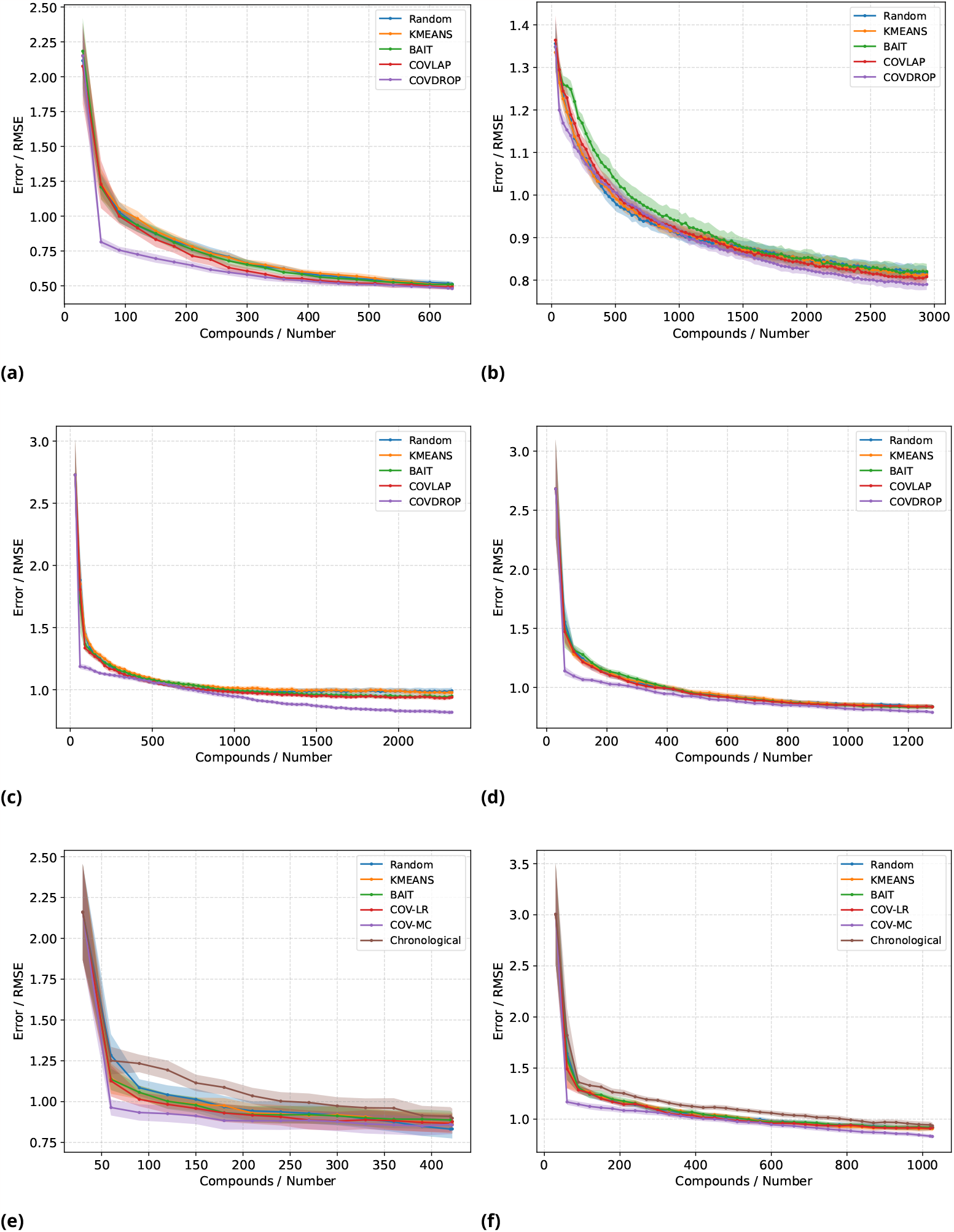
Performance of different methods for optimizing models for ADMET and affinity related datasets and batch selection strategies: a) the cell effective permeability b) lipophilicity. Panels c–f are affinity measurements to various targets proteins: c) GSK3β (ChEMBL) d) MMP3 (ChEMBL) e) Renin (Sanofi-Aventis) f) FXa (Sanofi-Aventis)

The overall shape of the RMSE profiles is impacted by the statistics of the target values in each dataset. For instance, for the plasma protein binding rate (PPBR) dataset, one can observe that all methods are suffering from large RMSE values early on. Specific to this case, there is an extreme imbalanced distribution for the target value of the source, as illustrated in Figure 8. Using a small number of compounds, the model gets a good insight of the most representative range, with a small peak following within the 300-400 samples, indicating a lack of training in underrepresented regions in early stages. In contrast to PPBR, hydration free energy (HFE) and effective cell permeability (Caco-2) the target distributions suffer less from skewness and spreadiness, as shown in Figure 8. For HFE and Caco-2 datasets COVDROP is the clear winner reaching RMSE within 10% of the final RMSE after testing only 36 and 450 compounds, respectively.

We also tested much larger datasets. For example, the aqueous solubility dataset (AS) (Figure 1 a) is significantly larger than those presented in Figure 2. There we see much slower convergence of RMSE values which we attribute to the normal distribution of the target values (Figure 8) and larger size of the training data. This RMSE profile indicates that for the Solubility dataset, the BAIT method exhibits inferior performance compared to the other methods, while COVDROP demonstrates the smallest root mean square error (RMSE) on the full dataset compared to all other methods. Additionally, COVDROP outperforms all other methods starting at 400 compounds.

### Performance on affinity data

We next evaluated the methods on several affinity datasets from ChEMBL and Sanofi-Aventis with different protein targets. For each of these, a diverse set of ligands is screened for their affinity to a target protein, e.g. MMP-8. Figure 2c depicts the retrospective experiment for modeling the affinity to Glycogen synthase kinase-3β(GSK3β), and we observe a similar trend to the solubility experiments. COVDROP outperforms very early the other methods.

Figure 2d presents to the retrospective experiment predicting the activity of small molecules against Matrix-metalloprotease(ChEMBL) with COVDROP gain the best option.

The internal datasets provide the opportunity to compare batch selection methods with the actual order used to experimentally test the compounds. We observe that all batch selection methods (though not random) outperform the current human based ordering (Figure 2 e-f and 1 h-i). For instance, for Renin and MMP-8, COVDROP significantly outperforms the chronological batch selection by requiring 62 and 58 % less training data points.

Similar to the ChEMBL data, we observed that for most of the molecules tested within the Sanofi-Aventis datasets COVDROP significantly outperforms other methods. However, for this dataset we also observed very good performance of COVLAP on the FXa dataset (Figure 2f). Furthermore, the number of experiments to achieve the 10% higher than minimum RMSE threshold is 750, while other selection methods require at least 12% more experiments to obtain the same metric (Tabel 1). Figure 2e showcases the retrospective experiment on the Renin dataset. Similar to our previous observations, we observed that COVDROP significantly outperformed the other methods. Notably, the method exhibited a small root-mean-square error (RMSE) after the initial batch selection in comparison to the final RMSE attained, which was achieved by the other methods after over 200 experiments and multiple iterations.

To further quantify the improvement we present in Table 1 the number of experiments required by each method for reaching model with an error at most 10% higher than the minimum RMSE obtained using all the data across all selection methods. As the table shows, both of our methods outperform all other methods for most datasets, in some cases significantly so. For example, we observed that for smaller datasets COVDROP can improve the results very quickly leading to much better performance. For most of the datasets performance improvements are greater than 10% vs. Random selection. As mentioned before for PPBR dataset due to the imbalance nature of the data COVDROP underperforms and BAIT selection produces the beset result. Even when compared to the chronological order in which the internal compounds were tested (i.e. to the actual experimental cycle) we observe improvements of up to 62%. This holds true if we change the stopping criteria to 20% or 5% difference (Table 4 & 5).

## Methods

### Datasets

To benchmark the various batch selection methods, we have collected both private (Sanofi-owned) and public datasets that represent a diverse range of some of the most important molecular properties that scientists need to address when developing small-molecule drugs. We list below the benchmark datasets used in this work covering the properties related to the absorption and distribution pharmacokinetic processes. In addition to the ADMET related properties, the benchmark datasets include four Sanofi and six public datasets recording the affinity of small drug molecules to ten target proteins, such as kinase and GCPRs. Table 1 provides detailed information on all of these datasets.

### Affinity datasets

Affinity measures the strength of binding between the ligands and biological targets, such as proteins. It is a critical molecular property that determines the drug efficacy. Therefore, to validate our active learning strategy for building statistical models, several datasets from the public database ChEMBL ***Gaulton et al. (2012)*** and internal sources were used.

### ChEMBL datasets

To retrieve suitable datasets from ChEMBL (version 31, https://www.ebi.ac.uk/chembl/), a collection of six proteins representing multiple target families (Table 1, column: Class) was identified using Uniprot IDs and ChEMBL target IDs. These targets include the alpha-1a adrenergic and dopamine D2 receptors as members of the GPCR family, glycogen synthase kinase-3 beta (GSK3β) from the kinase family, Matrix-metalloprotease 3, also known as MMP3 or stromelysin as metalloprotease, the sodium channel Nav1.7 as ion channel and the peroxisome proliferator-activated receptor delta (PPARδ) as member of the nuclear hormone receptor (NHR) family. See Supporting Methods for details on how these datasets were derived.

### Sanofi datasets

Four internal sets with structure-activity relationship (SAR) data were used. These allow us to overcome limitations of public SAR datasets which may merge assay data from different sources and which do not provide specific information on the order in which experiments were carried out. Such information allows direct comparison of the active learning solution and current best practices. The first dataset comprises compounds for inhibiting serine protease factor Xa (FXa)***Nazaré et al. (2004); Nazare et al. (2012); Matter et al. (2002); Nazaré et al. (2005); Matter et al. (2005)***. The second dataset is for inhibiting the aspartyl protease Renin***Scheiper et al. (2010); Matter et al. (2011); Scheiper et al. (2011)*** The third dataset is for the Matrix-metalloprotease 8, MMP-8, (human neutrophil collagenase) and the final dataset provides is for agonists for the nuclear hormone receptor PPARδ (peroxisome proliferator-activated receptor δ).

Each data point in the datasets comprises the chemical structure of the molecule, represented as a Simplified Molecular Input Line Entry System (SMILES) ***Weininger (1988)***, and the target value, which denotes the molecular property as a scalar. Table 1 provides details on the target and size of the each of these datasets.

### Active learning for small molecule optimization

We use active learning to optimize experiment selection. We assume a setup whereby a user can select from among a pool of unlabelled samples to query for their labels, i.e., to test experimentally. Furthermore, batches of experiments are performed in rounds.

In a given round, we denote the set of possible samples to select for labelling as 𝒱, and the batch size as *B*. The *ideal* active learner will select a subset, ℬ ⊂ 𝒱, |ℬ | = *B*, of these experiments such that, after the experiments are performed and the labelled samples are added to the training data, upon averaging it is expected that the overall loss of the *resulting* models is minimized.

In the original active learning frameworks, only one sample is queried at a time ***Lewis and Gale (1994) Dagan and Engelson (1995)***. In this case, a straightforward and effective strategy is to query the sample on which the current model’s prediction has the highest epistemic uncertainty. While this works well for single queries, it is hard to use the same approach for batch active learning. In many cases, the most uncertain queries will be similar to each other, and so their uncertainties are not independent, and so choosing as a batch the *B* most uncertain samples will be redundant, and not the most effective use of the queries ***Azimi et al. (2012)***.

In this paper we tested several different methods for selecting batches, including two new methods we developed. These two new methods are similar to “select the least certain samples” in spirit, selecting batches which maximize the total covariance of their uncertainty (estimated in different ways).

### Problem Formulation: Batch Active Learning for Deep Learning Regression Models

We consider a batch active learning scenario with multiple rounds of selection, {1, 2, …, *T* }. At the *t*-th round, let ℒ^(t)^ be the labeled dataset and 𝒱 ^(t)^ be the unlabeled dataset. To be clear, 𝒱 ^(t)^ and ℒ^(t)^ are determined only after the selection method has had a chance to examine 𝒱 ^(*t*−1)^ and ℒ ^(*t*−1)^ .

Since in our analysis we usually consider the selection problem in a *single round*, we omit superscripts ^(*t*)^where it won’t cause confusion.

Let *f*_*θ*_ *∶* 𝒳 → ℝ be the trained model, determined by its parameter *θ* ∈ Θ. As usual for regression problems, *θ* is chosen to approximately solve the following optimization problem,

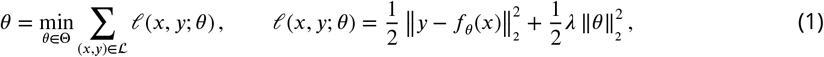

where 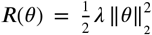 is the *L*^2^-regularization term for the weights *θ*. The optimization problem is (approximately) solved with a variation of SGD (in our case the popular ADAM***Kingma and Ba (2014)***), by iterating over multiple mini batches from the training set ℒ.

In Active Learning, a *selection method S* is a function

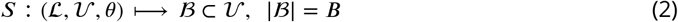

That is, *B* samples are selected from the unlabeled dataset 𝒱 by the active learning algorithm, and the selection may use the information of the given sets 𝒱 and ℒ, as well as the current supervised model *f*_*θ*_, which is trained using the labeled dataset ℒ.

Typically, selection is done by (approximately) solving an optimization problem,

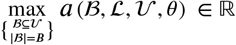

where the *acquisition function, a*, depends on the selection method. In a single selection round, ℒ, 𝒱 and *θ* are given to the selector, and from the selector’s point of view are fixed; therefore, in this case *a* depends only on the choice ℬ.

### Predictive uncertainty

Estimating predictive uncertainty is essential to our batch selection methods. Predictive uncertainty can be broadly divided into: *epistemic* uncertainty and *aleatoric* uncertainty, which may be roughly understood as *model uncertainty* and *noise*, respectively.

Aleatoric uncertainty is predictive uncertainty due to the inability of the model to give a precise prediction, even when the model parameters are uniquely specified. If *θ* are the model parameters, then we denote this distribution *p*_*Al*_ (*y*|*x, θ*).

Epistemic uncertainty is predictive uncertainty due to uncertainty in the model’s parameters. This is precisely the uncertainty which could be reduced by more labelled data, and therefore is the relevant quantity for active learning. In the strict Bayesian setting, if *f*_*θ*_ *∶* 𝒳 → ℝ is the model corresponding to parameters *θ*, and *p*(*θ*) is the posterior distribution given the training data 𝒟, this distribution is

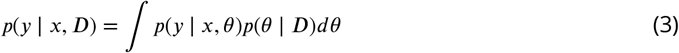

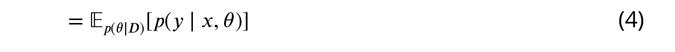

Calculating this distribution exactly is intractable, and so we use approximations, described in Section 16.

### Batch selection via determinant maximization

In the batch setting, selecting only those points with the epistemic uncertainty may lead to a batch in which uncertainties are highly correlated, and thus wasted experiments. I.e., if a sample *x* has the highest estimated variance, then small perturbations of *x* will have similarly high variance, and a variance-only selection method will give a high score to this batch of similar (and thus correlated) samples.

We thus seek to select the batch with the highest total independent uncertainty. This can be computed by selecting the batch with the highest joint entropy, which, under the assumption of a normal distribution, is the highest log-determinant of the (epistemic) covariance.

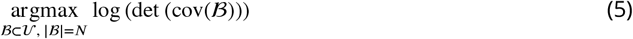

Even if we already know the epistemic covariance for any batch (a problem we address in the next section), finding the strict maximum of our acquisition function is NP-hard ***Ohsaka (2022)***. Therefore we use an approximate maximization technique: We randomly generate a collection of batches as starting points, each containing *N* distinct samples independently chosen from a distribution proportional to the quantile of the variances. Then we select the best *M* < *N* of these batches as starting points for optimization. Then, for each starting point, we optimize the batch element-wise, i.e., changing the first element to optimize the covariance, then changing the second, and so on, doing several passes until the process reaches equilibrium. Then, we select the highest-scoring final batch.

The log-determinant is computed using the Cholesky decomposition, which is a bit better than 𝒪(*N*^3^). However, the optimization step (substituting a new point into a batch one at a time) is just a rank-1 update to the batch’s covariance, and there are *O*(*N*^2^) rank-1 updates to the Cholesky decomposition, which we use.

#### Algorithm

Batch optimization

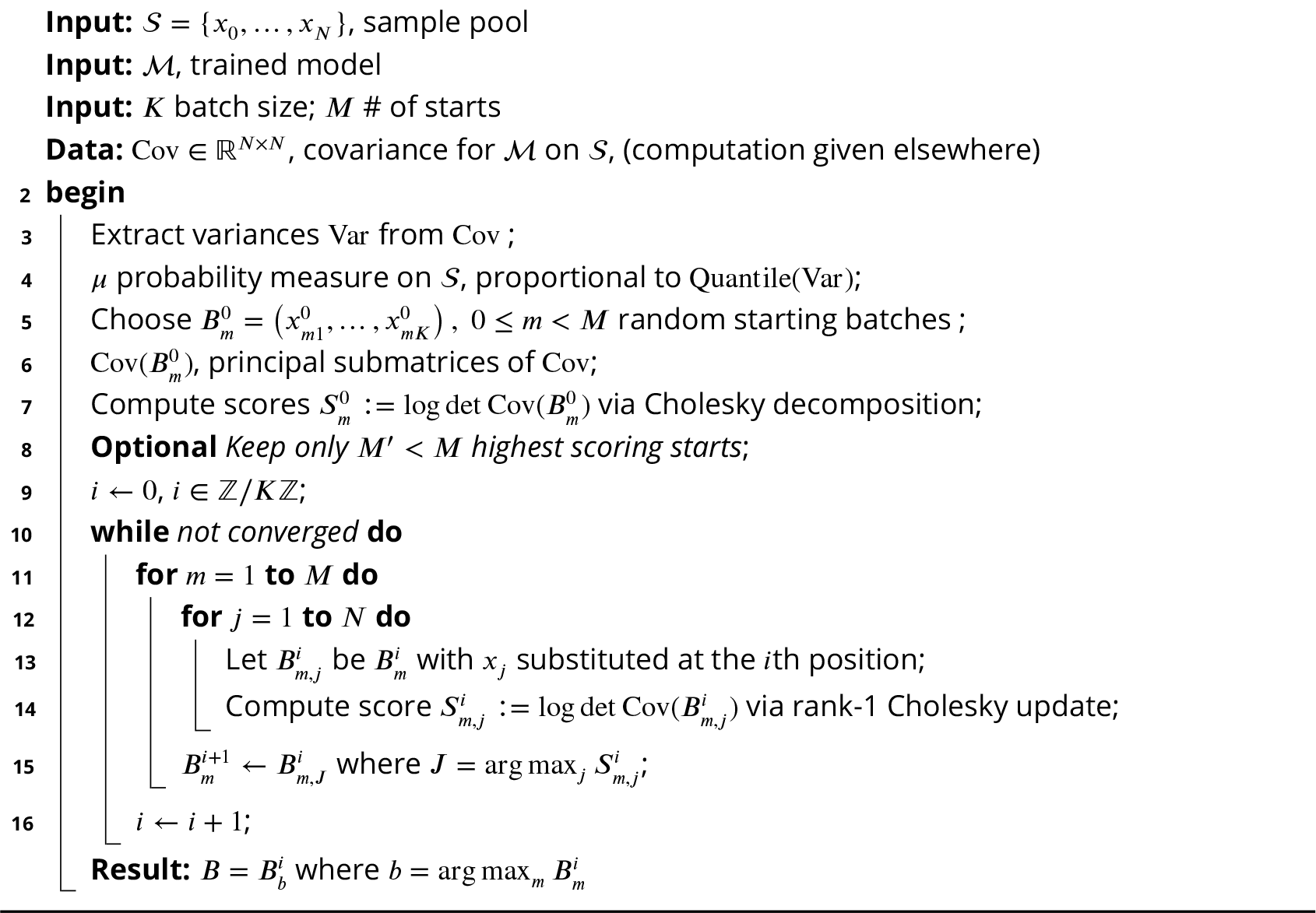

This leaves open the question of how to approximate predictive posterior of the model, and hence its covariance matrix over the sample pool. We address this in the following sections.

### Approximation of the posterior distribution

A straightforward way to get a distribution of predictions is to train an ensemble of models ***Lakshminarayanan et al. (2017)***, and to use the outputs of these models to give an ensemble of predictions. However, this approach involves multiple rounds of retraining, which is resource intensive. Furthermore, this ensemble of models will not sample from the Bayesian posterior, and there is no guarantee that it will be diverse.

Instead, we take a more economical approach by leveraging only one trained model. The idea behind this method is that the optimal parameters of a trained deep regression model, i.e. *θ*^∗^, are the *maximum a posteriori* (MAP) estimation of an equivalent Bayesian deep regression problem.

Thus, the Bayesian inference of the posterior of *θ* is approximated by leveraging the computed MAP estimation of *θ*^∗^. As it is shown in Eq. (1), the optimal *θ*^∗^ could be treated as the MAP estimation of the probabilistic model of {*Y, θ*} where *Y* = {*y*}_*y*∈_ℒ.

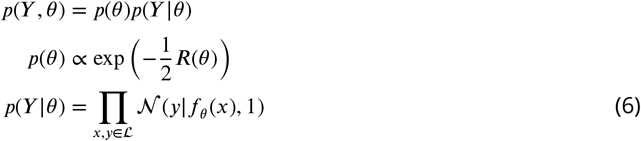

Two approximations for computing the epistemic covariance

We adapted two different methods to approximate the epistemic covariance:

- Monte Carlo Dropout

Monte Carlo dropout is a well-known technique for training neural nets. The usual approach is to turn off dropout during inference. However, if random dropout is applied during inference, then the model predictions will follow a probability distribution. It has been observed ***Gal and Ghahramani (2016)*** that this distribution is an approximation to the true Bayesian posterior. Therefore, we may sample an ensemble of predictions for a single sample, and use this distribution as a measure of the uncertainty of the prediction.

Dropout ***Hinton et al. (2012)*** is a well-known technique for better training of neural networks. With dropout, in the training stage, each of the neural network’s weights are randomly set to 0 with a certain probability (as defined by the drop out ratio, *r*). Previous work has demonstrated that models built with dropout are less prone to overfitting and that training with dropout is quite similar to variational inference of the probabilistic models ***Gal and Ghahramani (2016); Lawrence (2001)*** when assuming the following form of the posterior approximation *q*(*θ*),

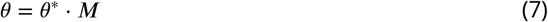

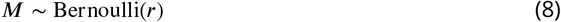

where *r* is the dropout ratio, *M*, the masks, are the {0, 1} variables of same shape as *θ*, and *θ*^∗^ are the weights obtained after training with dropout. We thus use dropout to obtain *S* predictions by sampling *S* masks {*M*_*s*_}_*s*∈1,2,⋯,*S*_ according to Eq. (8). We then use the new network to predict sample labels and construct the covariance matrix based on these predictions. Monte Carlo dropout can be defined as follows:

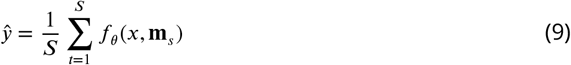

where ŷ is the predicted output of the model, *S* is the number of forward passes with dropout, *f*_*θ*_ is the model with parameters *θ, x* is the input to the model, and **m**_*s*_ is a binary mask that determines which units are dropped out during the *s*-th forward pass. In this equation, the model is run with dropout *S* times, and the predicted output is computed as the average of the outputs of each of these forward passes. This average is used as an estimate of the model’s predicted output and can be used to approximate the model’s uncertainty. Monte Carlo Dropout approach aligns with the principles of Bayesian Neural Networks but without the associated computational burden. Although this requires multiple forward passes during inference, the trade-off is beneficial, especially when understanding the uncertainty is paramount.

- Laplace Approximation

In addition to dropout, we also employed the *Laplace approximation* for estimating the posterior. This method assumes that the posterior of the model parameters is a multi-variate normal distribution centered at the MAP value, and with variance equal to the Fisher information, which can be found through a straightforward calculation. This simplifying assumption makes the approximation fast to use, as the computation relies only on MAP estimation and differentiation of the loss function.

We assume the following for the posterior probability distribution, *q*(*θ*):

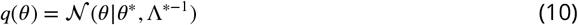

By matching 0^th^ and 2^nd^ derivatives, we find Λ, and thus our covariance matrix Λ^∗−1^. The covariance matrix Λ^∗−1^ does not have to be represented as a full matrix of shape *Q* × *Q*, which is usually very large for neural networks. For example, Laplace Redux ***Daxberger et al. (2021)*** further approximate the multi-variate normal distribution in Eq. (10) significantly reducing the computation cost. See Supporting Methods for additional details.

### Comparisons to other methods

To compare our proposed approach with prior approaches, for batch active learning we looked at three previous methods suggested for this task: Random selection ***Settles (2009)***, optimizing uncertainty and diversity based on information theory (the *BAIT* method), and an unsupervised method that is optimizing only for diversity ***Nguyen and Smeulders (2004)*** (a method we term *k-means sampling*). See Supporting Methods for details on these prior methods and how we used them here.

### Evaluation Experiments

For the retrospective experiment an existing labelled data set is selected to simulate an active learning experiment.

Thus we indirectly validated our posterior approximation by measuring the error with the respective method. The results from these retrospective experiments provided evidence for the accuracy and reliability of our approximation in representing the true posterior distribution.

We start by selecting a random subset (which is used as the initial set for all comparisons as well).

For our data the chemical structures in *X* are represented as molecular graphs using the `MolGraphConvFeaturizer` from the `DeepChem` library ***Kearnes et al. (2016)***. For each active learning cycle, models are trained using `DeepChem` ***dee (2016)***, a library which provides the implementation of the Graph Convolutional Neural Networks (GCNN) for molecular systems ***Kipf and Welling (2016)***.

For accuracy as well as model performance we use the root mean squared error (RMSE). See Supporting Methods for details on how the RMSE is computed.

### Neural network architecture

For all our datasets we use the “neural fingerprints” class of models described in ***Duvenaud et al. (2015)***, as implemented in the DeepChem library GraphConvModel ***contributors (2022)***. See Supporting Methods for complete details. Although our method is compatible with, PyTorch, TensorFlow, and Keras frameworks, the benchmarks presented here are performed using the Keras framework within Deepchem suite. In order to enforce deterministic behaviour of the models for each selection methods and active learning rounds, during the training the model weights were manually loaded at the beginning of each active learning loop. We perform multiple runs of the retrospective experiments for each methods.

## Code and Data availability

Code and all data sets used in this study are available as a python package at Sanofi-Public GitHub page. The package is called `ALIEN` (`Active Learning in data Exploration`) and can be downloaded: https://github.com/Sanofi-Public/Alien

