## Supplement for "Deep Batch Active Learning for Drug Discovery"

[1]R&D Data & Computational Science, Sanofi, , Cambridge, MA, United States; [2]Digital Data & Technology, Industriepark Höchst 65926 Frankfurt Germany; [3]Synthetic Molecular Design, Integrated Drug Discovery, Sanofi-Aventis Deutschland GmbH, Industriepark Höchst, Building G838, 65926 Frankfurt am Main, Germany; [4]Molecular Design Sciences, Integrated Drug Discovery, Sanofi, Vitry-sur-Seine, 94403, France; [5]Digital Data, Sanofi, Shanghai, China

[†]M.B.,R.L.,H.M., G.H.,C.G., Z.B. and S.J. designed research; M.B., S.M., R.L., A.C.N.,S.J., H.M., P.M., L.K.A., and S.R. performed research; M.B., S.M., A.C.N.,S.R., C. G., G.H. , H.M., P.M., and A.K. analyzed data; M.B., S.M., A.C.N., S.J., and Z.B. wrote the paper

### Supplement

#### Related work

**scikit-activeml**: This tool is a widely used machine learning library for Python that includes several active learning algorithms, such as uncertainty sampling, query by committee, and expected error reduction*Kottke et al.* (*2021*).

**ALiPy**: this active learning library is built on the scikit-learn package. It provides a unified interface for using active learning in scikit-learn and supports both stream-based and batch-based active learning *Tang et al.* (*2019*).

**ModAL**: This library provides a range of active learning algorithms, such as uncertainty sampling and query by disagreement. It also includes tools for evaluating the performance of active learning algorithms*Danka and Horvath* (*2018*).

#### Methods

##### Covariance Computation for the Last-Layer Laplace Approximation

In 1D regression tasks, the last layer is a linear map from the last-layer embedding space to the outcome,

$$x \longmapsto \theta \cdot x$$

Without loss of generality, we assume we have a linear model. Using the Laplace approximation, we compute a covariance matrix for the parameters $\theta$:

$$\Sigma_\theta = \left(\nabla^2_\theta \mathcal{L}(D,\theta)\right)^{-1}|_{\theta_{MAP}} \quad (1)$$

$$\nabla^2_\theta \mathcal{L}(D,\theta)|_{\theta_{MAP}} = \lambda^{-2} I + \sigma^{-2} \sum_{i=1}^{N} x_i x_i^T, \quad (2)$$

where $\lambda$ is the $L_2$ regularization constant and $\sigma$ is the aleatoric uncertainty. Up to a constant scaling of $\Sigma_\theta$, we may set $\sigma = 1$. In fact, a typical choice of prior uses $\lambda = 1$ as well.

Given the covariances $\Sigma_\theta$, we may compute covariances between $f_\theta(x_i)$ and $f_\theta(x_j)$, for our linear model $f_\theta$:

$$\Sigma_f = x_i^T \Sigma_\theta x_j \quad (3)$$

This calculation may be applied directly to the last layer of our neural nets, where the $x_i$'s are now the last layer embeddings and $\theta$ are only the last layer weights.

In our implementation, we are severely constrained by the fact that we need covariances, not just variances. We only approximate the posterior for weights in the last layer ($\boldsymbol{w}$), which is has a simple form. Weights for the other layers are approximated as completely certain at their MAP values. Previous works ***Daxberger et al.*** (***2021***) demonstrated that such a simplification still leads to uncertainty quantification which is adequate for many tasks.

### Batch Active learning via Information MaTrices (BAIT)

Batch Active Learning via Information MaTrices (BAIT) ***Ash et al.*** (***2021***) is a recent method which uses gradient embeddings combined with Fisher information to determine the optimal batch for selection. BAIT tries to select a batch, $\mathcal{B}$, which has the highest mutual information with the model weights, i.e., a batch whose uncertainty is collectively correlated most strongly with the uncertainty in the model weights. This is operationalized by (the trace of) the *Fisher matrix*.

In the case of a Bayesian linear regression model, BAIT provably minimizes the Bayes Risk, which represents the difference between the ground truth weights, $\boldsymbol{w}^*$, of the "true" model and the estimated weight, $\hat{\boldsymbol{w}}$. So BAIT is a generalization of this provably-optimal strategy to nonlinear models, where it is no longer optimal, but sometimes performs well.

One of the main challenges of BAIT is that one cannot afford to compute the Fisher information over all the weights of a deep regression model, so, like many selection strategies, it approximates the method by computing the Fisher matrix only for the last layer weights.

It can be proved that the Bayes risk after acquisition of batch $\mathcal{B}$ can be represented as follows,

$$\begin{aligned} \mathrm{BayesRisk}(\mathcal{B}) &= \sigma^2 \mathrm{tr}(J_{\mathcal{B}}^{-1} J_{\mathcal{U}}) \\ J_{\mathcal{B}} &= \sum_{z\in\mathcal{B}} \boldsymbol{z}\boldsymbol{z}^\top + \lambda\sigma^2 I \\ J_{\mathcal{U}} &= \sum_{z\in\mathcal{U}} \boldsymbol{z}\boldsymbol{z}^\top \end{aligned} \tag{4}$$

where $\lambda$ and $\sigma^2$ are hyper parameters of the Bayesian Linear Regression model. Note that the Bayes risk has nothing to do with the labels of the samples in $\mathcal{B}$.

For neural networks, a set $\mathcal{B} \subset \mathcal{U}$ of $B$ samples is chosen as follows:

$$\mathcal{B}^* = \underset{\mathcal{B}\subset\mathcal{U}}{\mathrm{argmin}}\, \mathrm{tr}\left(\left(\sum_{x\in\mathcal{B}} I(x;\phi)\right)^{-1}\left(\sum_{x\in\mathcal{U}} I(x;\phi)\right)\right) \tag{5}$$

where $I(x;\phi)$ is the Fisher matrix defined as the expectation over the current model $p(y|x,\phi)$ before acquisition as follows,

$$I(x;\phi) = \mathbb{E}_{p(y|x,\phi)} \nabla^2_{\phi} \ell(x,y;\phi) \tag{6}$$

It is straightforward to see that for Bayesian linear regression model, the Fisher information matrix for a single sample $\boldsymbol{z}$ is proportional to $\boldsymbol{z}\boldsymbol{z}^\top$, thus the objective in Eq.(5) approximately generalize the simple linear regression in Eq. (4)

### $k$-means Sampling

For focusing only on diversity, we relied on Zhdanov ***Zhdanov*** (***2019***) and used $k$-means clustering to partition the unlabeled dataset $\mathcal{U}$ into a defined number of centroids $\mathcal{K}$ with the size $k$, where $k = B$ and $\mathcal{B} \subset \mathcal{U}$, and then selecting a representative sample to be labeled as $\mathcal{B}$ ***Nguyen and Smeulders*** (***2004***) where

$$\mathcal{B} = \mathrm{argmin}_{z\in\mathcal{U} c\in\mathcal{K}} \|\boldsymbol{z} - c\|_2^2 \tag{7}$$

The L2 norm is minimized between the last layer embedding $\boldsymbol{z}$ and centroids $c$ where $c \in \mathcal{K}$. The algorithm works by iteratively using the $k$-means clustering algorithm to cluster the unlabeled examples in $\mathcal{U}$ into $k$ clusters, selecting the next nearest neighbor to its centroid $c$(cf. Eq. (7)) and adding it to the labeled data set $\mathcal{L}$.

### Metrics for performance evaluation

RMSe is used to asses the method performance. RMSE is a measure of the average squared difference between the predicted values and the true values. It is calculated as:

$$RMSE = \sqrt{\frac{1}{N}\sum_{i=1}^{N}(y_i - \hat{y}_i)^2}$$

where $y_i$ is the true value for the $i^{th}$ data point in the test set, and $\hat{y}_i$ is the predicted value for the $i^{th}$ data point.

*pearson* correlation on the test set which can be defined as follows:

$$r = \frac{\sum_{i=1}^{n}(x_i - \bar{x})(y_i - \bar{y})}{\sqrt{\sum_{i=1}^{n}(x_i - \bar{x})^2}\sqrt{\sum_{i=1}^{n}(y_i - \bar{y})^2}}$$

where $x_i$ and $y_i$ are the predicted values and true values for the test set, respectively, and $\bar{x}$ and $\bar{y}$ are the means of the predicted values and true values, respectively.

### Neural network architecture

Our convolutional network used 2 layers of width 128, with a final pooling layer of width 16, before a linear output to a single real value. The learning rate and dropout probability of were set to 0.001 and 0.05, respectively. The chosen training batch size is based on the dataset size, normally ranging between 32 and 64 with the objective of smoothing the loss curves. For each selection method, 25 active learning cycles/runs were executed, with an initial randomized set of 30 samples and a batch of selected candidates of size 32.

Due to the small size of some of the datasets used in this study, it was necessary to account for the overfitting. Therefore, we used an early stopping protocol where the model performance on a validation set is continuously monitored, and training is stopped once it has gone for 100 epochs without improvement of the best-performing model, as defined by the validation metric, i.e., MSE. The datasets were split into train, validation, and test sets, with a ratio of 70%, 15%, and 15% of the dataset, respectively.

### Chemical Frameworks for Active Learning

`ChemML` and `DeepChem` are python libraries for applying machine learning methods to problems in the field of chemistry. Both libraries were developed to provide a high-level interface for working with complex datasets and models in these domains, and it includes a wide range of tools and functionality. However, they lack support for state-of-the-art deep batch active learning.

**ChemML**: This package provides limited tools for active learning of regression models using expected model change (EMC) and query by committee (QBC) methods ***Haghighatlari et al.*** (***2019***).

**DeepChem**: this module provide variety of tools for chemical data management, featurization, and machine learning models that can be interfaced with widely used machine learning frameworks.

### Results

Table 1 presents the details associated with each dataset used for method evaluation in this strudy. To curate the affinity data originated from ChemBL database, all compound entries associated to a target ID were extracted from ChEMBL (column: Retrieved Cpds), while only those with existing $IC_{50}$ or Ki values in ChEMBL were kept. For PPARδ compounds with an $EC_{50}$ value to characterize agonistic activity, were kept instead. For all other targets, antagonistic or inhibitory activity was reported. As data in ChEMBL are compiled combining different literature sources, they summarize assay data from different laboratories. While those data might not be fully comparable due to differing assay setups and protocols, the standardized ChEMBL $pIC_{50}$ value (pChEMBL) is used as first attempt to compare molecules across different literature sources. However, we are aware of the limitations of this comparison across series with assay data from different laboratories. All datasets were filtered by existing pChEMBL values (column: Filtered Cpds), converted from Smiles to SDF format using Corina (version 2021), and structurally validated using RDKit (version 2022.09, https://www.rdkit.org/). Identical structures not considering stereochemistry were merged and the most active pChEMBL value was used to represent this entry (column: Final Dataset). These datasets are then used for building statistical models.

### Internal Data

Furthermore, four internal sets with structure-activity relationship (SAR) data were collected for this validation study. Our goal is to overcome limitations of public SAR datasets merging assay data from different sources and to provide information on registration dates for evaluating time series.

During the data collection for the new affinity data, provided by Sanofi-Aventis, for each compound, the minimum submission date, i.e., the first time, a batch for this compound was registered in our dataset, is converted into an integer to indicate the month of registration. This integer exists for all datasets and defines time series for active learning based on series progression after a certain time, as usual in industrial lead optimization settings.

The first dataset comprises 1468 compounds as inhibitors of the serine protease factor Xa (FXa)***Nazaré et al.*** (***2004***); ***Nazare et al.*** (***2012***); ***Matter et al.*** (***2002a***); ***Nazaré et al.*** (***2005***); ***Matter et al.*** (***2005***) representing multiple internal series. Those series include amino-acids, aminoquinolines, azole-carboxamides, benzamidines, carboxybenzamides, indole-carboxamides, ketopiperazines, orthobenzamides, oxybenzamides, piperidine-benzamides, pyrrolidines and aminoethanesulfonamides (taurines). The inhibition of human factor Xa is given as pKi value (negative logarithm of inhibition constant Ki) from a standardized and harmonized internal biochemical assay.

The second dataset is for the aspartyl protease Renin***Scheiper et al.*** (***2010***); ***Matter et al.*** (***2011***); ***Scheiper et al.*** (***2011***) and contains 604 inhibitors from multiple series including acylguanidines, benzhydryls, benzylpiperidines, indole-carboxamides, azaindole-carboxamidines, indolizine-carboxamides, pyrazole-carboxamides and spiropiperidines. The inhibition of human Renin is provided as $pIC_{50}$ value (negative logarithm of $IC_{50}$) from a standardized and harmonized internal biochemical assay. Procedure of the assay were carried out as described in ***Ostrem et al.*** (***1998***).

The third dataset is for the Matrix-metalloprotease 8, MMP-8, (human neutrophil collagenase)***Matter et al.*** (***2002b***); ***Matter and Schudok*** (***2004***); ***Matter and Schwab*** (***2000***, 1999); ***Matter et al.*** (***1999***) with 652 inhibitors from multiple series, which includes amino-acids, dialkyl-acids, pyrimidine-bisamides, tetrahydroisoquinolines and thiophene-carboxylates. Inhibition of human MMP-8 is given as $pIC_{50}$ value from a standardized and harmonized internal assay.

For the execution of the assay, Biological activities were tested in a luminescence spectrometer using a specific quenched fluorigenic substrate. The tests were carried out in 96-well plates. Each well had a buffer, enzyme, and drug. The reaction was initiated by adding the substrate after a 15-minute preincubation. The enzymatic reaction speed was measured without and with various ligand concentrations. All measurements were conducted twice for accuracy. The results analyzed the ligand-dependent inhibition, and IC50 values were determined using standard software.***Matter et al.*** (***2002b***); ***Matter and Schudok*** (***2004***); ***Matter and Schwab*** (***2000***, 1999); ***Matter et al.*** (***1999***)

The final dataset provides 878 partial and full agonists for the nuclear hormone receptor PPARδ (peroxisome proliferator-activated receptor delta) from multiple series including aryloxyacetic acids, arylsulfonamides, carboxylates, oxadiazolones, thiadiazoles and trifluoromethyl-sulfonamides***Keil et al.*** (***2011***).

These four datasets were otherwise prepared in analogy to the public datasets from ChEMBL before subjecting them for building statistical SAR models.

Further characterizations of the internal data sets can be found in Figure 8.

### ADMET dataset

- **Lipophilicity**: Lipophilicity that reflects a drug propensity in a lipid environment, e.g. fatty tissue, impacts the drug metabolism and uptake. In this dataset the partition coefficient of 4,200 drug molecules between octanol and water at pH 7.4 is provided.
- **Aqueous Solubility**: The water solubility of a molecule impacts its biological activity, such as bioavailablity and target binding. Molecular structure and the type of functional groups on the drug molecule affects its interaction with water and solubility. In this dataset the LogS, where S is the molar concentration of 9,982 molecules are available along with their 2D structural descriptors, e.i. SMILES are reported.
- **Caco-2 (Cell Effective Permeability)**: Permeability of the cell membrane to a drug impacts drug absorption, thereby, cell permeability is an important ADMET property. in this dataset the Caco-2 cells which are a cell line derived from human colon cancer cells, are used in vitro for estimating the drug permeability coefficients. in this dtaset the permeability coefficient for 906 drug molecules are included.
- **Hydration Free Energy**: As a energy equivalent of Aqueous solubility, hydration free energy impacts ADMET properties of the drug candidate. In this dataset, 642 free energy values derived from experimental and computational methods are provided.
- **Plasma Protein Binding Rate (PPBR)**: The extent ot the binding of drug molecules to plasma proteins contributes to their cell uptake. For instance, drug molecules with weak binding to globulins render better tissue distribution and liver metabolism. Thus, PPBR value prediction is essential for *in silico* drug design. In this data set the PPBR for 1,614 molecules are obtained from a multiple spices which lead to a heterogeneous set.

| Target protein | Target class | # Compounds | category | source |
|---|---|---|---|---|
| FXa | Protease | 1468 | affinity (pKi) | Sanofi-Aventis |
| PPARδ | Nuclear receptor | 878 | affinity($pEC_{50}$) | Sanofi-Aventis |
| MMP-8 | Metalloprotease | 652 | affinity($pIC_{50}$) | Sanofi-Aventis |
| Renin | Angiotensinogenase | 604 | affinity($pIC_{50}$) | Sanofi-Aventis |
| A1AR | GCPR | 1362 | affinity(pChEMBL) | ChEMBL |
| D2 receptor | GCPR | 6853 | affinity(pChEMBL) | ChEMBL |
| GSK3β | Kinase | 3323 | affinity(pChEMBL) | ChEMBL |
| MMP3 | Metalloprotease | 1830 | affinity(pChEMBL) | ChEMBL |
| Nav1.7 | Ion channel | 5722 | affinity(pChEMBL) | ChEMBL |
| PPARδ | Nuclear receptor | 1257 | affinity(pChEMBL) | ChEMBL |
| - | Lipophilicity | 4200 | ADMET | PyTDC, Ref: ***Wenlock and Tomkinson*** (***2015***) |
| - | Caco-2 (Cell Permeability) | 906 | ADMET | PyTDC, Ref: ***Wang et al.*** (***2016***) |
| - | Hydration Free Energy | 642 | ADMET | PyTDC, Ref: ***Wu et al.*** (***2018***) |
| - | Solubility | 9982 | ADMET | PyTDC,Ref: ***Sorkun et al.*** (***2019***) |
| - | Plasma Protein Binding Rate | 1640 | ADMET | PyTDC, Ref: ***Wenlock and Tomkinson*** (***2015***) |

**Table 1.** The collection of the datasets with their corresponding target protein and property used for method evaluation. The chemical structure of the compound is represented using SMILES notation, which allows its computational analysis. The affinity value is given in pChEMBL, a numerical value that ranges from 0 to 1 and which provides an estimate of the likelihood that a particular compound will bind to its target. For the ADMET category the process that the drug undergoes within a living organism are named based on the molecular property aimed to be predicted. Each dataset is split into training, validation, and test with 0.7,0.15,0.15 ratio.

Figure 1 shows the method evaluation of the 9 datasets from ADMET and affinity related properties, such as aqueous solubility, hydration free energy, and plasma protein binding rate, PPARδ. For instance, panel (f) depicts the retrospective experiment predicting the affinity of a small molecule to the Alpha-1a adrenergic receptor. Our analysis revealed that the COVDROP method significantly outperformed all other tested methods, including COVLAP, by achieving a 20% lower root-mean-square error (RMSE) value with only 270 compounds, whereas $k$-means and random selection methods required 390 compounds to attain the 20% higher than minimum calculated RMSE value.

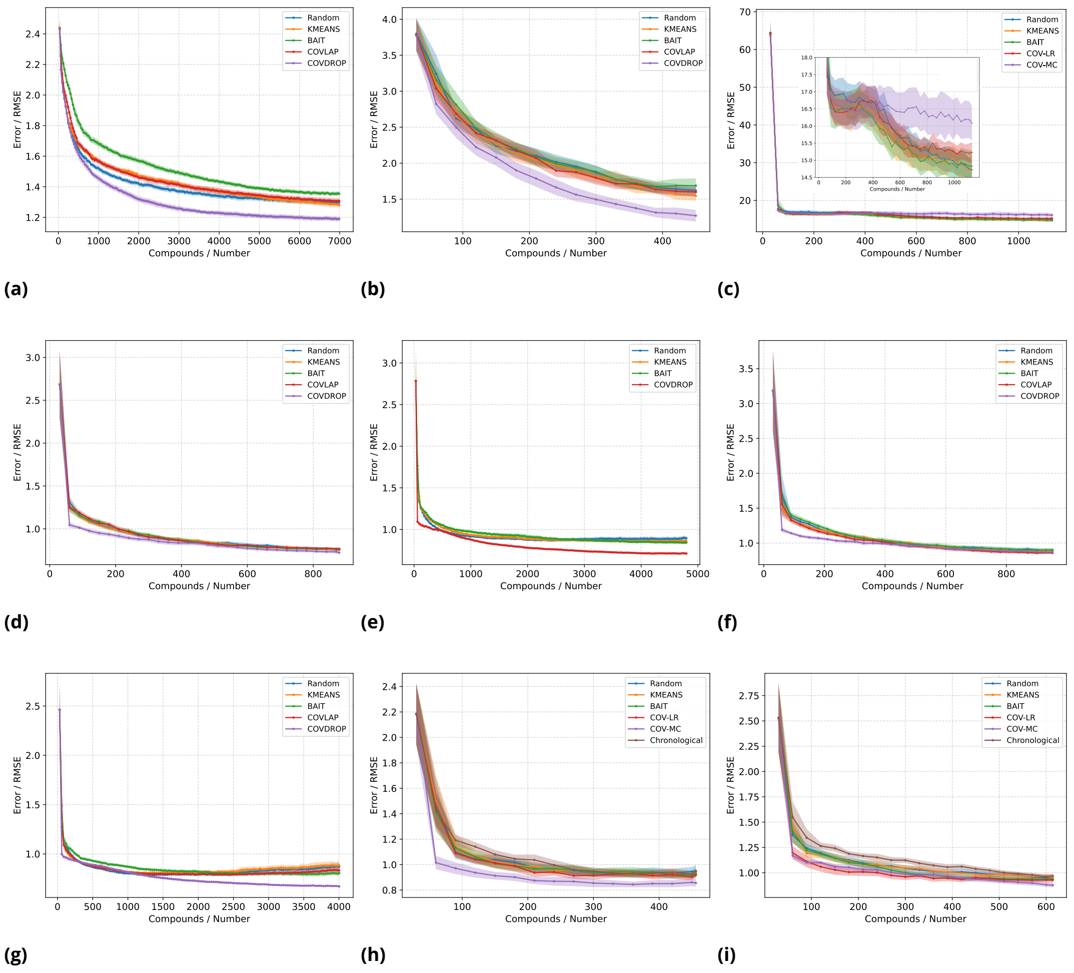


**Figure 1.** Retrospective experiments with ADMET and affinity related datasets and batch selection strategies. ADMET: a) Solubility b) hydration free energy c) plasma protein binding rate. Affinity to various target proteins: d) Peroxisome proliferator-activated receptor delta (PPARδ: ChEMBL) e) Dopamin D2 receptor f) Alpha-1a adrenergic receptor g) Sodium channel protein type IX alpha subunit h) Matrix-metalloprotease 8 i) Peroxisome proliferator-activated receptor delta (PPARδ: Sanofi-Aventis)

---

Interestingly, COVLAP started outperforming COVDROP by reaching the model error with a 10% higher than the minimum calculated RMSE value with the complete training set, as well as for the 5% mark, requiring only 690 compounds to achieve this benchmark. This is the first instance where our second proposed method has also outperformed COVDROP. panel(g) displays the retrospective experiment on the Sodium channel protein type IX alpha subunit dataset. Initially, COVDROP outperforms the other methods, but after 230 compounds, it catches up with all the other methods. It is worth noting that BAIT does not perform well, showing a higher RMSE compared to all other curves.

Similarly, in the MMP-8 dataset (Sanofi-Aventis) (Figure 1h) COVDROP outperforms every other selection method, as it only requires 90 experiments to reach 20% of the minimum obtained RMSE (Table 4), whereas others need 3 to 4 times more experiments to get to the same point. Lastly, we have the retrospective experiment conducted for the PPARδ dataset (Sanofi-Aventis) illustrated in Figure 1i). Here, we observe once again an outstanding performance from COVDROP, with only 210 experiments needed to achieve the threshold. Surprisingly, an even better performance for COVLAP shows up, with the lowest RMSE most of the experimental time.

To illustrate the efficiency of our selection method algorithms with respect to the Random se-

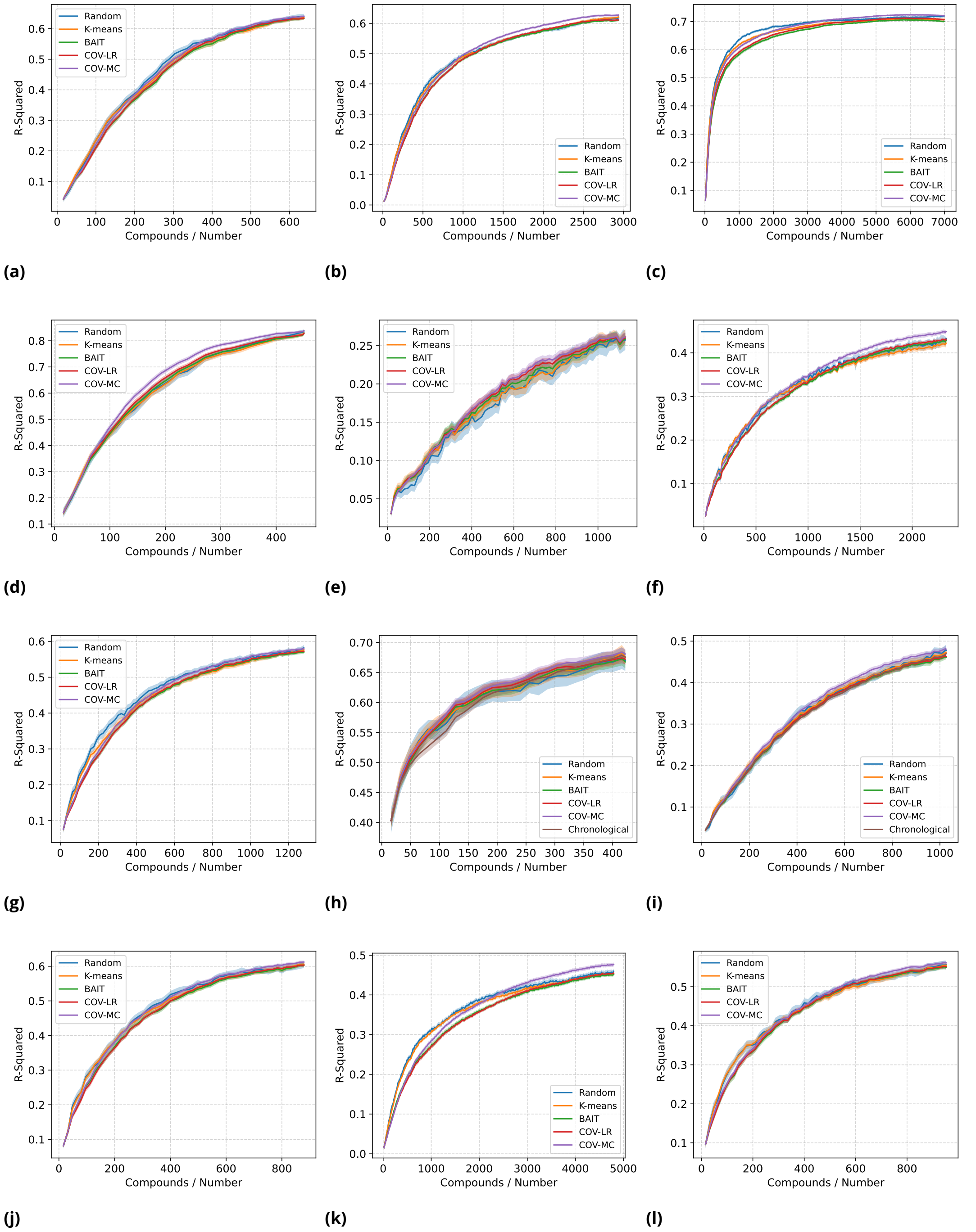


**Figure 2.** R-Squared plots for retrospective experiments with ADMET and affinity related datasets and batch selection strategies. ADMET: a) the cell effective permeability b) lipophilicity c) solubility d) hydration free energy e) plasma protein binding rate. Affinity to various target proteins: f) GSK3β (ChEMBL) g) MMP3 (ChEMBL) h) Renin (Sanofi-Aventis) i) FXa (Sanofi-Aventis) j) Peroxisome proliferator-activated receptor delta (PPARδ: ChEMBL) k) Dopamin D2 receptor l) Alpha-1a adrenergic receptor

---

lection, we performed the computation time analysis of these two selection methods on the GPU and CPU processors. The MMP-8 data set was used in this analysis, a 2-layer deep GCN model was trained for 1 active learning round. As it is shown in Table 2, our selection methods is about twice more expensive than the random batch selection on a traditional computer with 64 CPUs while on

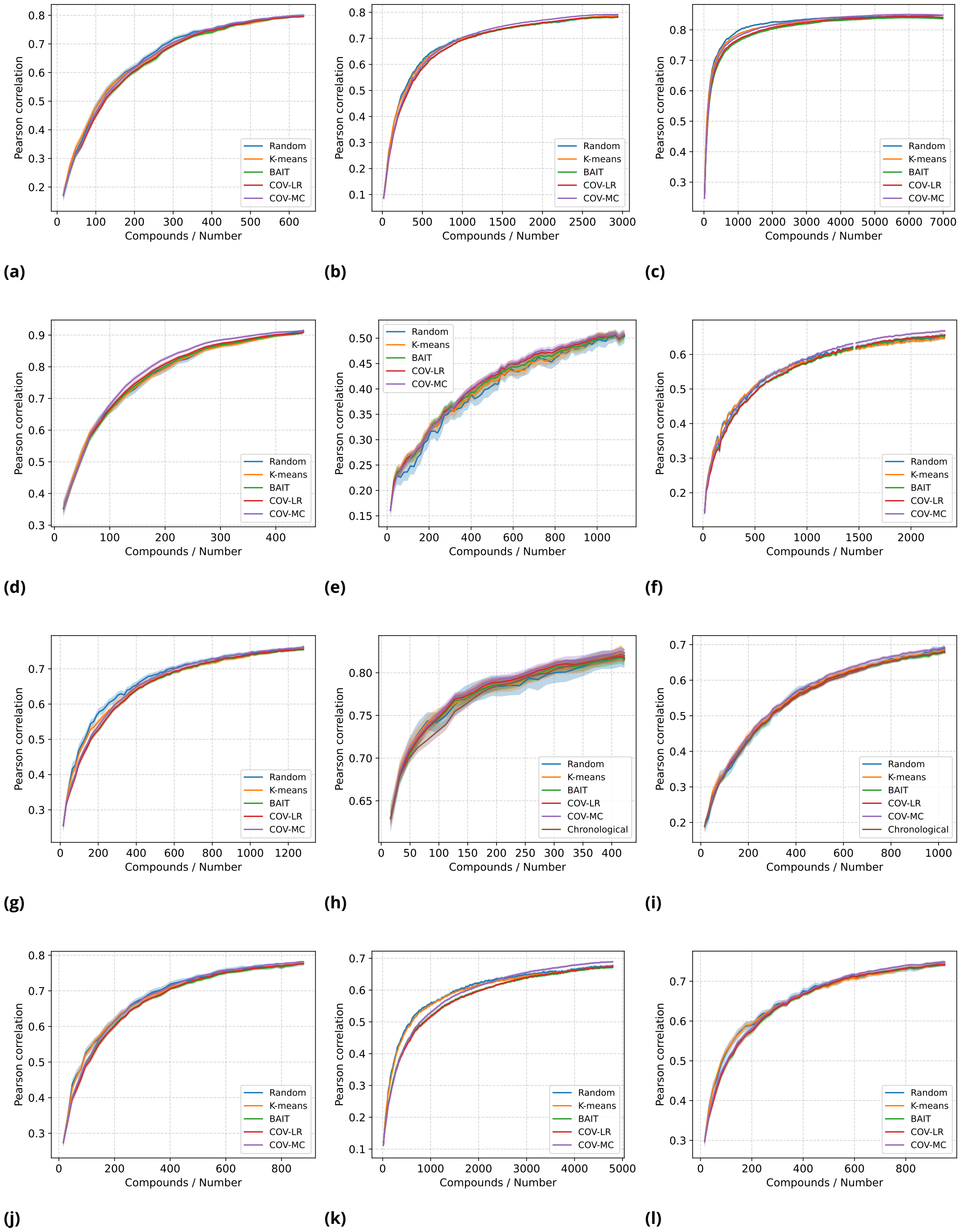


**Figure 3.** Pearson correlation coefficient plots for retrospective experiments with ADMET and affinity related datasets and batch selection strategies. ADMET: a) the cell effective permeability b) lipophilicity c) solubility d) hydration free energy e) plasma protein binding rate. Affinity to various target proteins: f) GSK3β (ChEMBL) g) MMP3 (ChEMBL) h) Renin (Sanofi-Aventis) i) FXa (Sanofi-Aventis) j) Peroxisome proliferator-activated receptor delta (PPARδ: ChEMBL) k) Dopamin D2 receptor l) Alpha-1a adrenergic receptor

---

a GPU our method is 35% more expensive.

Figure 6 demonstrates the scatter plots of the $k$-means clustered datasets. Table 3 presents an evaluation of these clusters using Silhouette coefficients***Rousseeuw*** (***1987***), where a higher score indicates better-defined clusters. Our observation shows that the performance of $k$-means selector

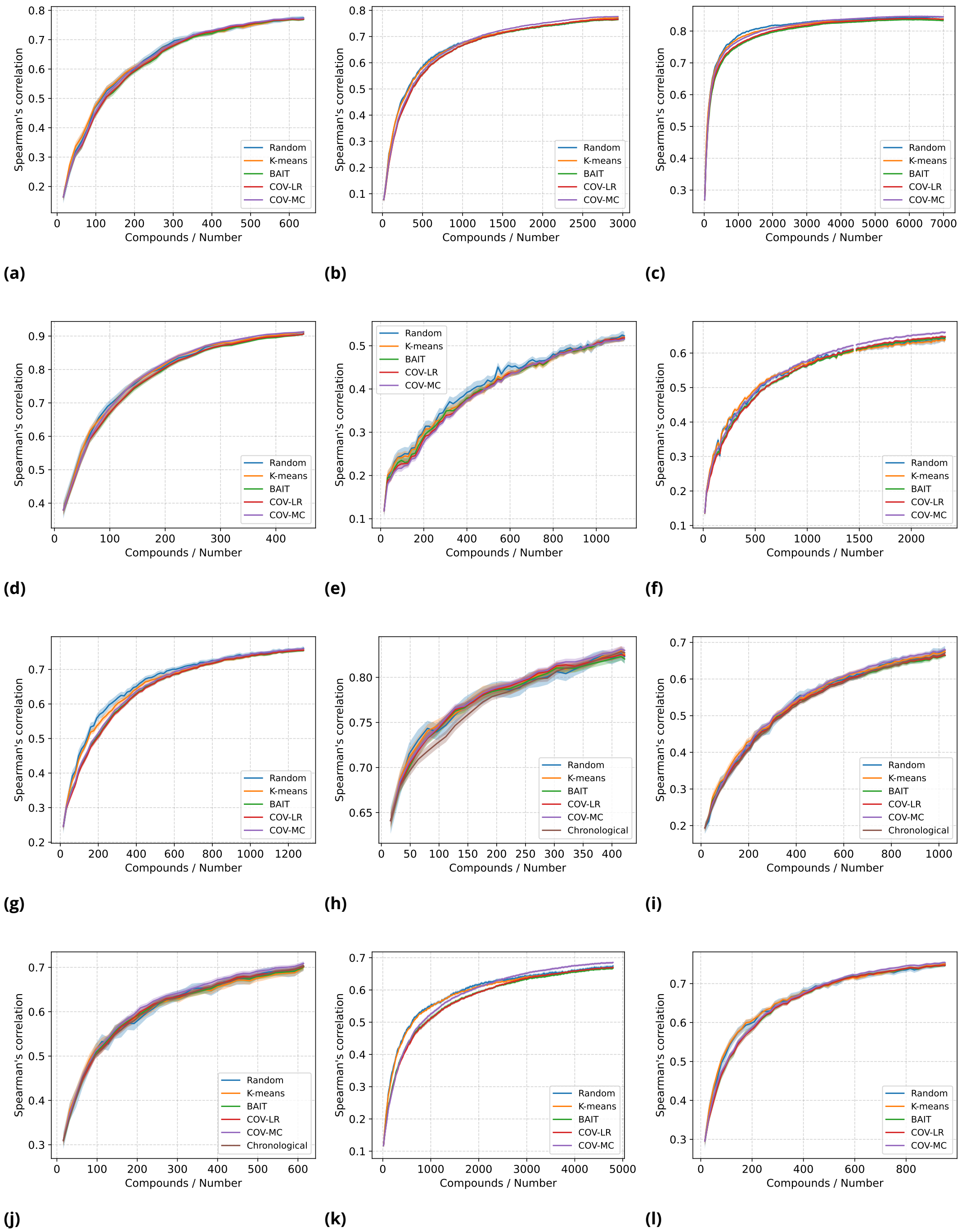


**Figure 4.** Spearman's correlation plots for retrospective experiments with ADMET and affinity related datasets and batch selection strategies. ADMET: a) the cell effective permeability b) lipophilicity c) solubility d) hydration free energy e) plasma protein binding rate. Affinity to various target proteins: f) GSK3β (ChEMBL) g) MMP3 (ChEMBL) h) Renin (Sanofi-Aventis) i) FXa (Sanofi-Aventis) j) Peroxisome proliferator-activated receptor delta (PPARδ: ChEMBL) k) Dopamin D2 receptor l) Alpha-1a adrenergic receptor

---

is inversely correlated with the Silhouette coefficient. Specifically, the $k$-means selector performed best on the plasma protein binding rate dataset, which had the lowest Silhouette coefficient. However, it underperformed in comparison to other methods on the lipophilicity and solubility datasets, which had better defined clusters with high Silhouette coefficients which may point towards exis-

| processor | information | Random | COVDROP |
|---|---|---|---|
| GPU | 1 × Tesla V100-SXM2-16GB | 21.92 | 29.55 |
| CPU | 1 × Intel(R) Xeon(R) @ 2.50GHz | 10.82 | 19.52 |

**Table 2.** Timing (minutes) comparison of batch selection methods on CPU and GPU processors for completing an active learning round of experiment using the same model architecture as described in the Method section.

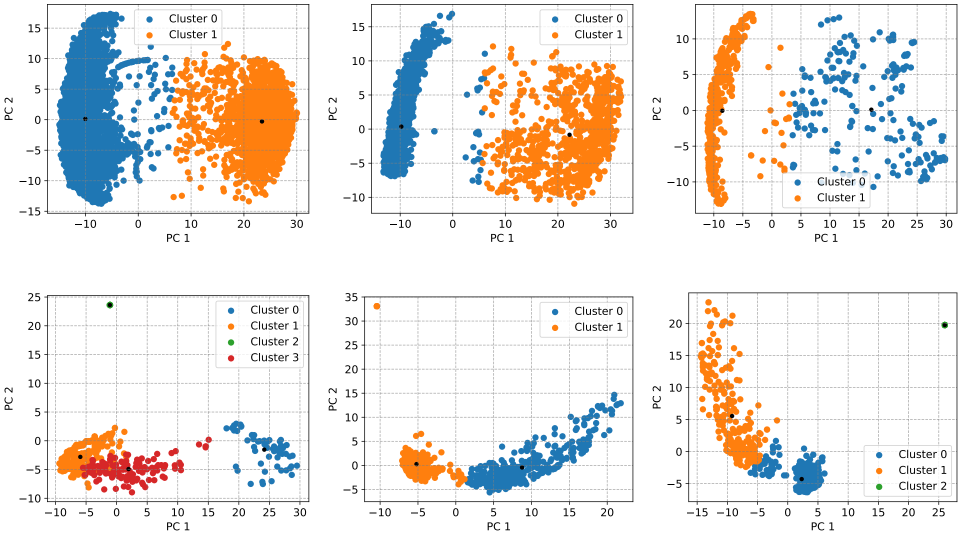


**Figure 5.** $k$-means clustering with varius ADMET related datasets using GCN embeddings: a) Solubility b) Lipophilicity c) the cell effective permeability d) Hydration Free Energy e) the Plasma Protein Binding Rate f) the Clearance-hepatocyte

| Dataset | Silhouette Coefficient |
|---|---|
| Solubility | 0.4635 |
| Lipophilicity | 0.5425 |
| Caco-2 | 0.4275 |
| Hydration Free Energy | 0.4309 |
| Plasma Protein Binding Rate | 0.3135 |

**Table 3.** Silhouette coefficients for clustered datasets

tence of significant number of outliers in d–f.

Figure 7 illustrates the effect of number of the greedy optimization rounds of the batch selection on the Covariance method. The results show that the number of greedy optimization rounds slightly impacts the model performance. The solubility dataset was utilized monitor the effect of this parameter on the model performance.

Tables 4 & 5 show the performance and batch selection methods on evaluation datasets on reaching the RMSE values at 20 and 5 % higher than minimum RMSE values.

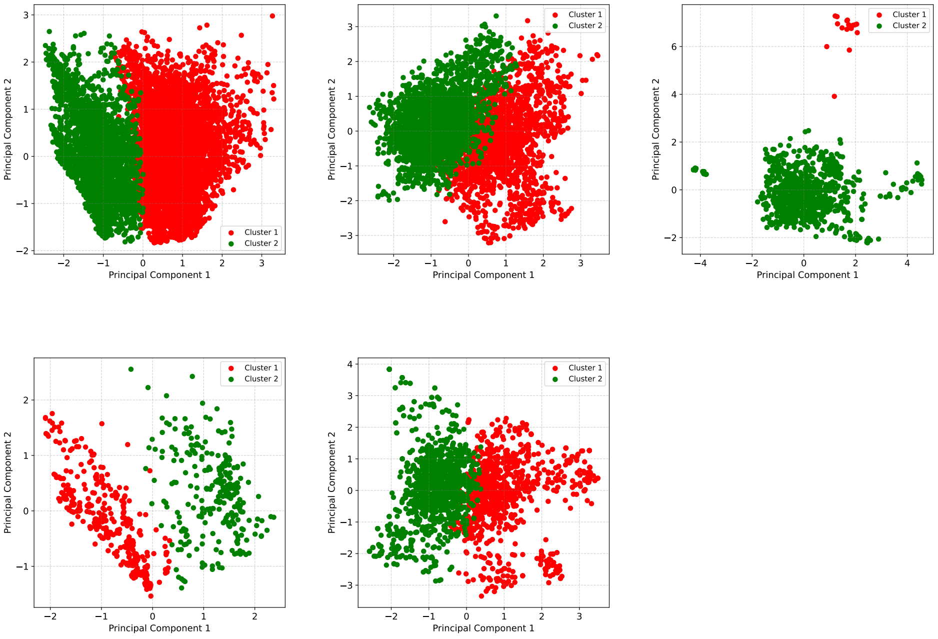


**Figure 6.** $k$-means clustering with varius ADMET related datasets using circular fingerprints with radius of 3 Åand Bit size of 1024: a) Solubility b) Lipophilicity c) the cell effective permeability d) Hydration Free Energy e) the Plasma Protein Binding Rate f) the Clearance-hepatocyte

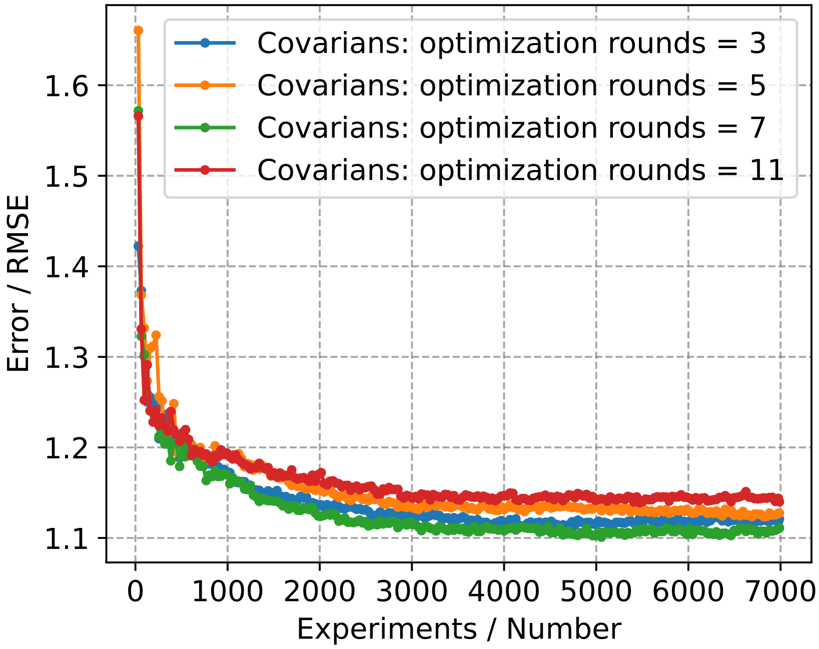


**Figure 7.** fine tuning number of the greedy optimization sounds in the Covariance selection method. the solubility dataset was used to monitor the dependency of the Covariance selection method on the number of optimization rounds.

| Dataset | COVDROP | COVLAP | $k$-means | BAIT | Random | Chron. | NC | % gain |
|---|---|---|---|---|---|---|---|---|
| Caco2 | 330 | 360 | 480 | 420 | 420 | - | 637 | 21.4 |
| Lipophilicity | 810 | 780 | 750 | 930 | 690 | - | 2940 | -17.4 |
| HFE | 300 | 450 | 450 | 450 | 450 | - | 450 | 33.3 |
| Solubility | 1170 | 2700 | 2880 | 4140 | 1860 | - | 6988 | 37.1 |
| PPBR | 60 | 60 | 60 | 90 | 60 | - | 1130 | 0.0 |
| GSK3β | 840 | 990 | 2130 | 1170 | 1230 | - | 2325 | 31.7 |
| MMP3 | 420 | 480 | 570 | 510 | 510 | - | 1280 | 17.6 |
| PPARδ | 330 | 390 | 420 | 420 | 420 | - | 879 | 21.4 |
| A1AR | 270 | 390 | 420 | 420 | 390 | - | 952 | 30.8 |
| Nav17 | 1080 | 1140 | 1650 | 2370 | 960 | - | 4004 | -12.5 |
| D2 | 1170 | 3840 | 4797 | 3840 | 4797 | - | 4797 | 75.6 |
| S Renin | 60 | 120 | 150 | 150 | 180 | 270 | 422 | 66.7 |
| FXa | 480 | 480 | 570 | 540 | 600 | 810 | 1026 | 20.0 |
| MMP-8 | 90 | 180 | 180 | 150 | 210 | 240 | 456 | 57.1 |
| PPARδ | 210 | 150 | 300 | 270 | 300 | 450 | 614 | 30.0 |

**Table 4.** Number or required experiments for reaching model error with 20% higher than minimium calculated RMSE with the whole training set. The chronological based batch selection is represented under the Chron. columns

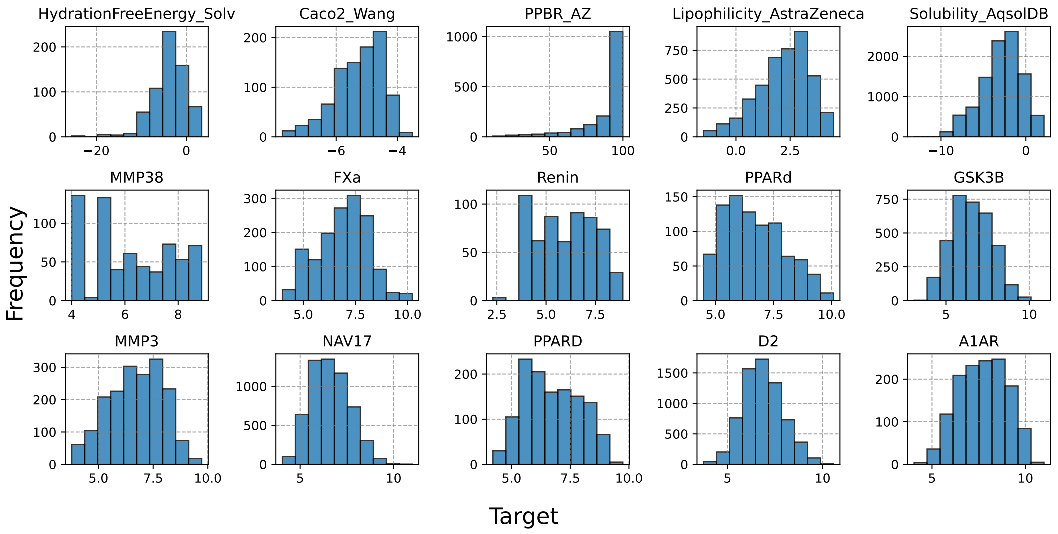


**Figure 8.** Distribution of target values for the different datasets from various sources used in this work. The datasets included in the figure are: HydrationFreeEnergy_Solv, Solubility_AqsolDB, Caco2_Wang, Lipophilicity_AstraZeneca, PPBR_AZ from Therapeutic Data Commons; FXa, PPARδ, MMP-8, and Renin from Sanofi's internal sources; and A1AR, D2, GSK3β, MMP3, Nav17, and PPARδfrom CHEMBL.

| Dataset | COVDROP | COVLAP | $k$-means | BAIT | Random | Chron. | NC | % gain |
|---|---|---|---|---|---|---|---|---|
| Caco2 | 540 | 600 | 637 | 637 | 637 | - | 637 | 15.2 |
| Lipophilicity | 1890 | 2250 | 2430 | 2490 | 2520 | - | 2940 | 25.0 |
| HFE | 390 | 450 | 450 | 450 | 450 | - | 450 | 13.3 |
| Solubility | 3240 | 6988 | 6988 | 6988 | 6988 | - | 6988 | 53.6 |
| PPBR | 1130 | 660 | 600 | 540 | 690 | - | 1130 | -63.8 |
| GSK3β | 1620 | 2325 | 2325 | 2325 | 2325 | - | 2325 | 30.3 |
| MMP3 | 990 | 1280 | 1280 | 1280 | 1280 | - | 1280 | 22.7 |
| PPARδ | 660 | 879 | 879 | 879 | 870 | - | 879 | 24.1 |
| A1AR | 750 | 690 | 810 | 840 | 900 | - | 952 | 16.7 |
| Nav17 | 2550 | 4004 | 4004 | 4004 | 4004 | - | 4004 | 36.3 |
| D2 | 2850 | 4797 | 4797 | 4797 | 4797 | - | 4797 | 40.6 |
| Renin | 330 | 390 | 420 | 422 | 390 | 422 | 422 | 15.4 |
| FXa | 870 | 1026 | 1026 | 1026 | 1026 | 1026 | 1026 | 15.2 |
| MMP-8 | 210 | 456 | 456 | 456 | 456 | 456 | 456 | 53.9 |
| PPARδ | 510 | 614 | 614 | 614 | 614 | 614 | 614 | 16.9 |

**Table 5.** Number or required experiments for reaching model error with 5% higher than minimum calculated RMSE with the whole training set. . The chronological based batch selection is represented under the Chron. columns

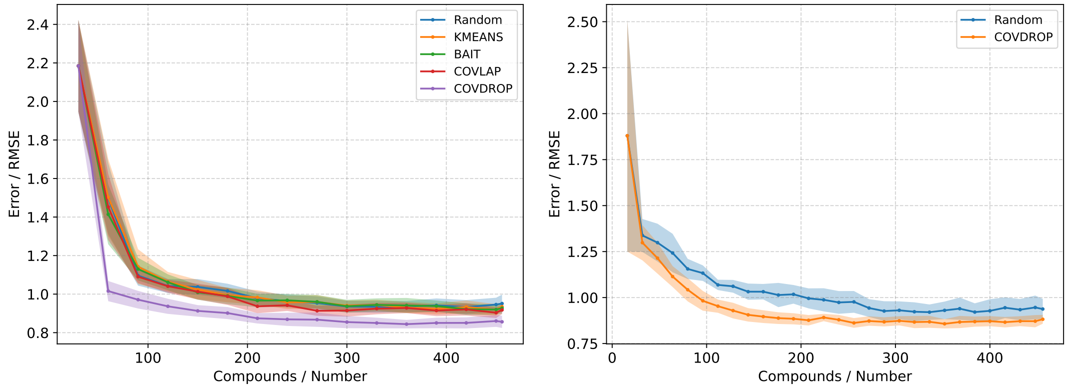


**(a)** **(b)**

**Figure 9.** the impact of model size on the RMSE profile. MMP-8 dataset is used in this test a) represents the larger model architecture. i.e. 2 GCN layers of 128 nodes each and batchsize of 32. b) small model architecture, i.e. 1 GCN layer of 16 nodes and batch size of 16.

Figure 9 depicts the impact of the model size on the RMSE profile.

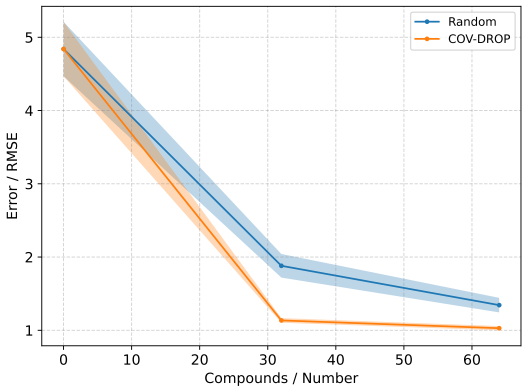


**Figure 10.** The selection performance of COVDROP vs. random selection, with empty initial set and an untrained model

### Toy example

We show a toy example with a synthetic dataset. The ground truth was chosen as a sinusoidal function on the unit square. However, the training set was sampled only from an elliptical subregion (Fig. 11). For our model, we used a simple 4-layer MLP, with width 128 and GeLU activation, and (of course) dropouts at rate .1.

After training the model on the subregion, we found good agreement between predictions and ground truth in the training region, and predictions getting steadily worse, the farther away from the training region (Fig. 12 **b**). We computed uncertainties (as std. dev.) using dropout during inference. We see that the uncertainty (Fig. 12 **d**) is highest outside of the training region, though there is not perfect agreement with the actual error (Fig. 12 **c**) since, as the saying goes, “a stopped clock is right twice a day.”

Finally, we ran the COVDROP algorithm to select a batch of 10 points for labelling, shown in Figure 14. The selected points tend to be in regions of high uncertainty, and/or at the edge the feature space.

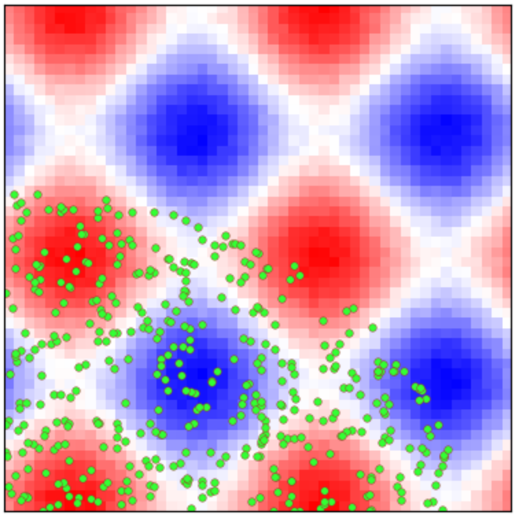

**Figure 11.** Ground truth function on unit square: red is positive, blue is negative; Green dots are the training set.

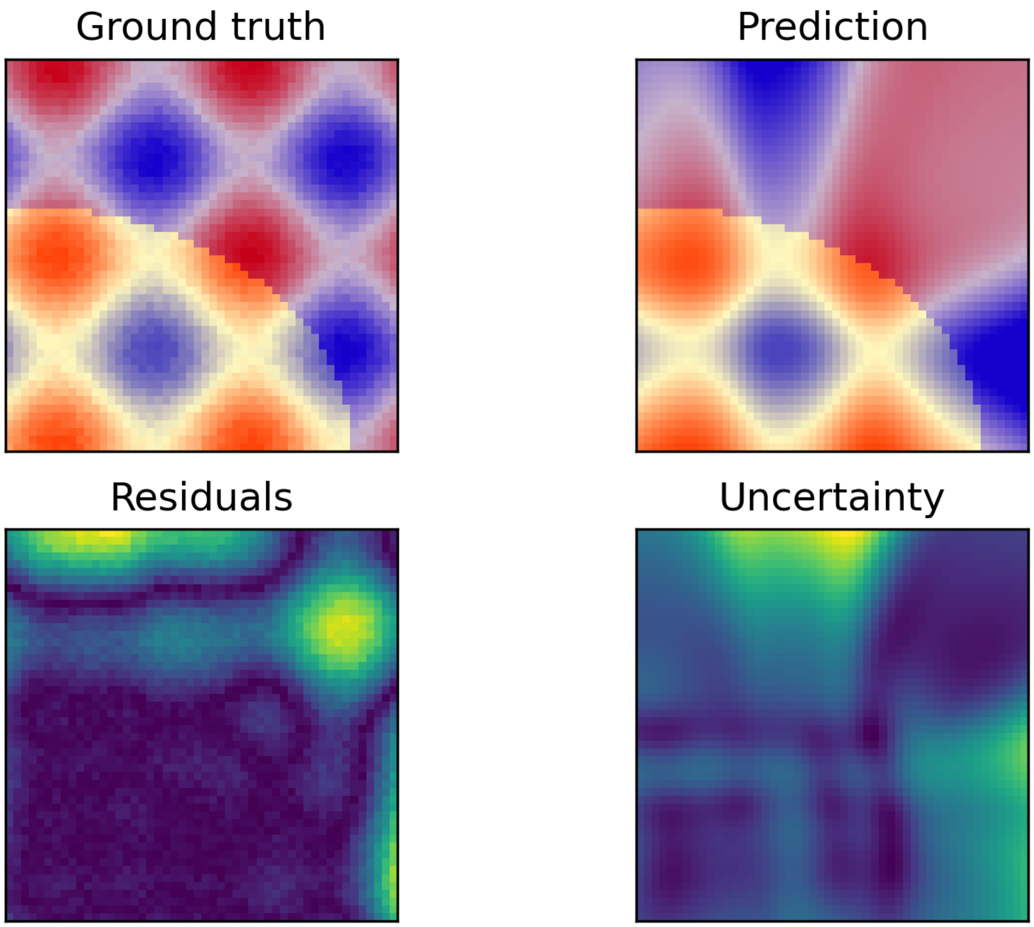


**Figure 12. a)** Ground truth (with training region highlighted). **b)** Trained model predictions. **c)** Absolute error. **d)** Model uncertainty.

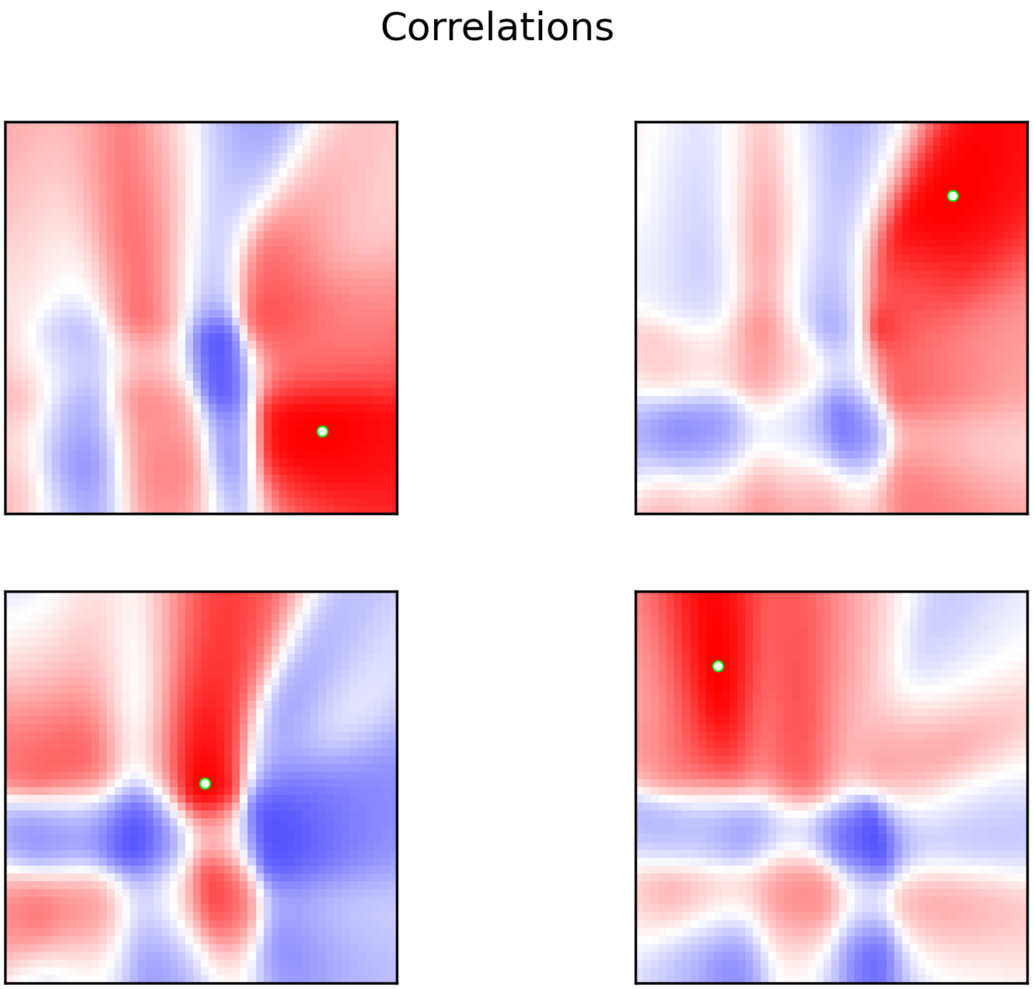


**Figure 13.** Posterior correlations between predicted values. Each square shows the correlation of one selected base point (highlighted white) with all the others. Red is positive, blue is negative.

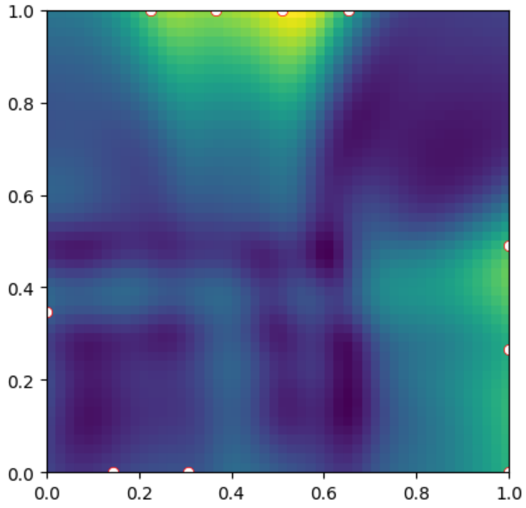


**Figure 14.** Batch of points for labelling (white dots), chosen
